# Spatial summation in the human fovea: the effect of optical aberrations and fixational eye movements

**DOI:** 10.1101/283119

**Authors:** William S. Tuten, Robert F. Cooper, Pavan Tiruveedhula, Alfredo Dubra, Austin Roorda, Nicolas P. Cottaris, David H. Brainard, Jessica I.W. Morgan

## Abstract

Psychophysical inferences about the neural mechanisms supporting spatial vision can be undermined by uncertainties introduced by optical aberrations and fixational eye movements, particularly in fovea where the neuronal grain of the visual system is fine. We examined the effect of these pre-neural factors on photopic spatial summation in the human fovea using a custom adaptive optics scanning light ophthalmoscope that provided control over optical aberrations and retinal stimulus motion. Consistent with previous results, Ricco’s area of complete summation encompassed multiple photoreceptors when measured with ordinary amounts of ocular aberrations and retinal stimulus motion. When both factors were minimized experimentally, summation areas were essentially unchanged, suggesting that foveal spatial summation is limited by post-receptoral neural pooling. We compared our behavioral data to predictions generated with a physiologically-inspired front-end model of the visual system, and were able to capture the shape of the summation curves obtained with and without pre-retinal factors using a single post-receptoral summing filter of fixed spatial extent. Given our data and modeling, neurons in the magnocellular visual pathway, such as parasol ganglion cells, provide a candidate neural correlate of Ricco’s area in the central fovea.

## Introduction

Vision science seeks to understand how the retinal image is encoded and processed by the visual system. A more specific question is how processing applied at each level of the visual pathways limits the information available to subsequent stages. One example is spatial pooling. Classically, spatial pooling has been investigated using the areal summation paradigm, in which an intensity-area reciprocity is observed at detection threshold for spatially uniform stimuli below a critical diameter (Ricco, 1877). This relationship is termed Ricco’s Law and implies that at detection threshold photons falling within an integration area are pooled completely by the visual system. Beyond Ricco’s area of complete summation, this relationship is no longer maintained and further increases in stimulus size yield smaller gains in sensitivity.

Ricco’s area of complete summation depends on a number of factors, including background intensity (Barlow, 1958; Glezer, 1965; Lelkens & Zuidema, 1983; Redmond, Zlatkova, Vassilev, Garway-Heath, & Anderson, 2013), the chromaticity and polarity of the stimulus and background (Brindley, 1954; Vassilev, Ivanov, Zlatkova, & Anderson, 2005; Vassilev, Mihaylova, Racheva, Zlatkova, & Anderson, 2003; Volbrecht, Shrago, Schefrin, & Werner, 2000), and distance from the fovea (Hallett, 1963; Inui, Mimura, & Kani, 1981; Khuu & Kalloniatis, 2015; Scholtes & Bouman, 1977; Wilson, 1970). Together, these results suggest that the summation area is shaped by the functional architecture of the post-receptoral visual pathways mediating stimulus detection at threshold. However, the neural underpinnings of spatial summation at threshold remain unclear, particularly in the fovea where the neuronal density of the retina and downstream circuitry is highest (Curcio & Allen, 1990; Curcio, Sloan, Kalina, & Hendrickson, 1990; Smallman, MacLeod, He, & Kentridge, 1996) and where pre-retinal factors such as optical aberrations and fixational eye movements inject spatiotemporal blur into the proximal stimulus that is difficult to disentangle from post-receptoral neural pooling.

To estimate the relative effect of pre-neural factors on foveal summation, Davila and Geisler (1991) compared behavioral data to summation curves generated by a computational observer that incorporated estimates of photon fluctuations in the stimulus, the optical properties of the ocular media, and the spatial arrangement and quantum efficiency of the photoreceptor lattice (Davila & Geisler, 1991). Their analysis suggested that foveal summation could be explained by optical factors without the need to posit post-receptoral summation. We reasoned that if there is essentially no post-receptoral summation at threshold for spot detection, then summation areas measured under aberration-free optical conditions should approach the dimensions of a single foveal cone (∼0.5 arcmin diameter).

More recently, foveal summation measurements were obtained using an adaptive optics (AO) vision simulator (Dalimier & Dainty, 2010). Despite correcting for ocular aberrations, this study yielded estimates of Ricco’s area in the fovea that were similar to those acquired previously with conventional stimulus-delivery platforms (reviewed in Davila and Geisler, 1991). However, Dalimier and Dainty did not compensate for fixational eye movements, and the use of a non-imaging AO system precluded objective confirmation of AO correction fidelity or stimulus focus onto the photoreceptor layer.

To unravel the relative contributions of high-order optical aberrations, fixational eye movements, and post-receptoral processes on spatial summation in the central fovea, we used a multi-channel adaptive optics scanning light ophthalmoscope (AOSLO) equipped with high-speed retinal tracking and stimulus delivery capabilities (Dubra & Sulai, 2011; Roorda et al., 2002; Yang, Arathorn, Tiruveedhula, Vogel, & Roorda, 2010). We found Ricco’s area to be essentially invariant to modest amounts of fixational eye motion and optical blur, suggesting that post-receptoral neural pooling plays an important role in spatial summation measured in the central fovea. Further, the summation areas we measured encompassed multiple foveal cones, more closely resembling the anatomical dimensions of parasol ganglion cell dendritic fields in the human fovea, suggesting that the magnocellular pathway mediates the detection of circularly-shaped increments at visual threshold (Swanson, Sun, Lee, & Cao, 2011; Volbrecht et al., 2000).

## Methods

### Retinal imaging and psychophysical testing with an AOSLO

We examined the effects of optical aberrations and fixational eye movements on photopic signal integration in the human fovea. Four subjects (one female, three male; age range: 29 to 56 years) with normal color vision and no known retinal pathology in the studied eye participated in the study. Prior to enrollment, informed consent was obtained from each subject. All subjects were experienced psychophysical observers and were aware of the purpose of the study. All study protocols adhered to the tenets of the Declaration of Helsinki and were approved by the Institutional Review Board at the University of Pennsylvania.

An AOSLO designed for multi-modal high-resolution retinal imaging (Dubra & Sulai, 2011; Dubra et al., 2011; Scoles et al., 2014) was modified to enable psychophysical testing. To achieve this, a field-programmable gate array (FPGA)-based image acquisition and stimulus control module was incorporated into the existing system architecture, facilitating the high-speed retinal tracking and light source modulation required for cone-targeted stimulus delivery (Arathorn et al., 2007; Yang et al., 2010). Analog signals from an H7422-50A photomultiplier module (PMT; Hamamatsu, Shizuoka Pref., Japan), positioned behind a confocal pinhole (1.13 Airy Disk diameters) to encode the instantaneous intensity of the focused infrared (λ = 795 nm) imaging beam, were mirrored and sent to both the FPGA acquisition module as well as to a separate frame grabber native to the existing AOSLO (HEL 2M QHAL E*, Matrox Electronic Systems Ltd, Dorval, Quebec, Canada). The FPGA-based acquisition system digitized the signals into 512-by-512 retinal images at 16 Hz using an analog-to-digital converter operating in coordination with h-sync and v-sync timing signals generated by the scanning control hardware. The sinusoidal distortion in pixel geometry introduced by the high-speed resonant scanner was measured by acquiring an image of a square calibration grid with 0.10 degree spacing; image frames were de-sinusoided in real-time using custom FPGA-based software. The resultant retinal videos enabled the extraction of retinal motion in real-time via a strip-based image registration (Vogel, Arathorn, Roorda, & Parker, 2006). The eye tracking signals were in turn used to control the timing of an acousto-optic modulator (AOM; Brimrose Corporation, Sparks, MD) capable of adjusting the intensity of the co-aligned stimulus beam (λ = 550 ± 15 nm; **Figure 1A**) at frequencies exceeding the 20 MHz pixel clock of the system (Poonja, Patel, Henry, & Roorda, 2005). The stimulus source was a supercontinuum laser (SuperK Extreme EXU-6 OCT, NKT Photonics, Birkerød, Denmark) whose peak wavelength and bandwidth were controlled by a tunable single-line filter (SuperK VARIA, ibid.).

**Figure 1.**
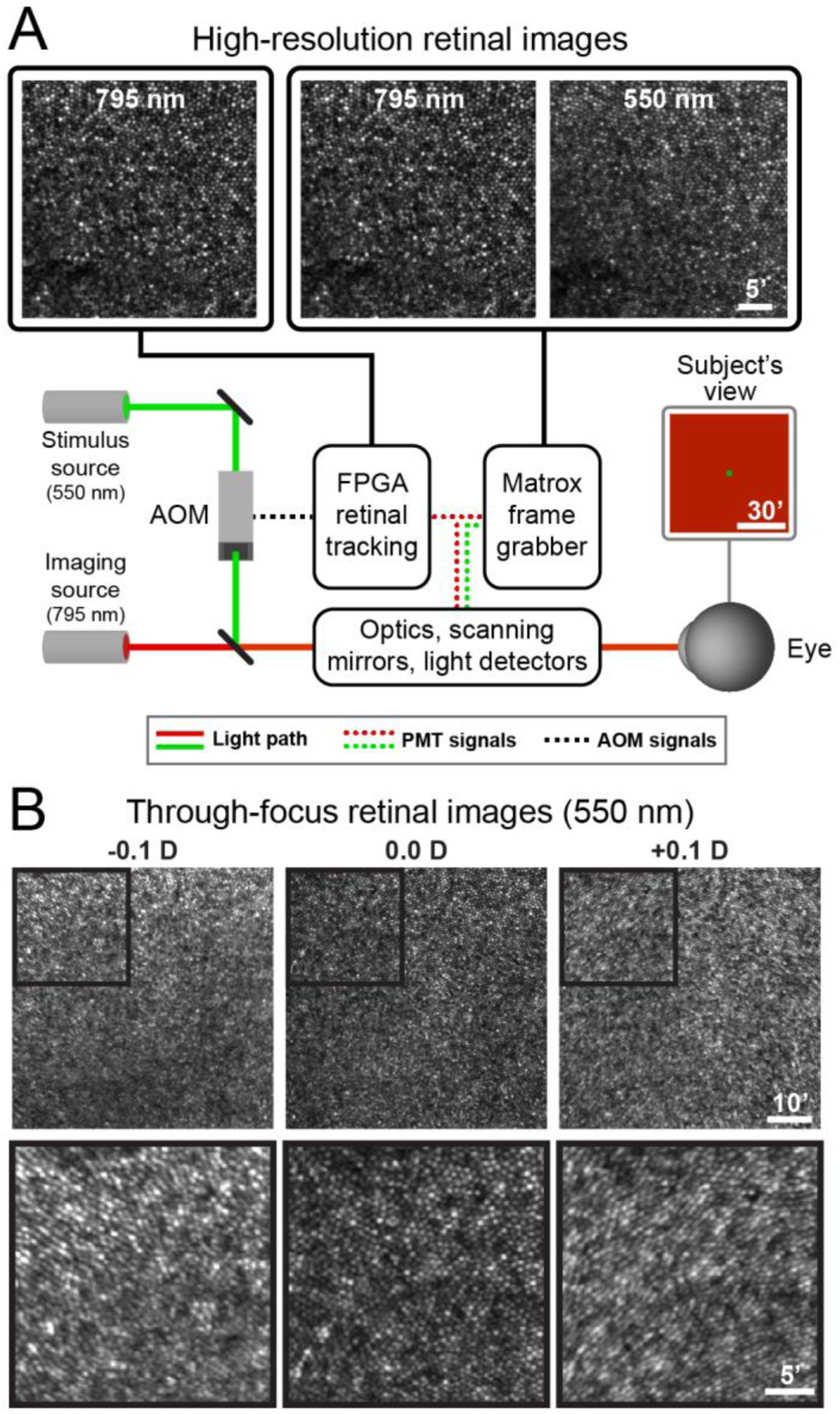
Features of the adaptive optics scanning light ophthalmoscope. (A) Schematic of the AOSLO used in this study. High-resolution retinal images could be acquired by digitizing signals from two independent imaging channels, each featuring a photomultiplier tube (PMT) positioned behind a confocal pinhole. The tightly-packed bright spots in the images in the upper left panels are individual cone photoreceptors near the subject’s fovea (bottom left corner). Each image was averaged from 40 registered video frames and cropped to 35×35 arcmin to highlight the cellular resolution of the AOSLO. The primary source for retinal imaging and eye tracking was a near-infrared superluminescent diode (795 nm); infrared PMT signals were sent to both the native frame grabber (for multichannel imaging) and a field-programmable gate array (FPGA) module (for real-time retinal tracking). The 795 nm image is duplicated in this schematic representation. A 550 nm image could also be acquired simultaneously with the 795 nm image via the native frame grabber. Stimulus patterns were delivered to the retina by modulating the 550 nm source with an acousto-optic modulator (AOM) controlled by the FPGA module. The subject viewed the 1.2 degree square imaging raster upon which circular increment stimuli were presented. See Methods for more details on imaging and psychophysical procedures. (B) The top row shows spatially-registered images of cone photoreceptors obtained with 550 nm light in the fovea of S2 across a range of focal depths; the fovea is near the center of each panel. Images were collected with prescribed amounts of defocus (in diopters, D; indicated by the text in each panel). All other aberrations were corrected by the deformable mirror. Best focus was determined subjectively by the examiner and assigned a value of zero diopters. Black squares outline regions presented at higher-magnification in the bottom row, where subtle image degradation is evident with small amounts of negative and positive defocus. Each image is averaged from 40 registered video frames.

The infrared PMT signals could be used to generate complementary retinal videos in the native frame grabber, which was also capable of digitizing signals from a second confocal channel (1.40 ADD pinhole) that permitted the simultaneous acquisition of full-frame images with the stimulation wavelength. The acquisition parameters of the native frame grabber were set to match those of the FPGA-based system as closely as possible, although videos acquired with this digitizer could not be de-sinusoided in real-time. To correct for any residual differences in image dimensions between the two systems, images of the calibration grid were collected simultaneously on the native (with sinusoidal distortion) and FPGA (without sinusoidal distortion) modules prior to a measurement session. The image transformation required to render the former at the pixel scaling of the FPGA-based images was derived using custom software in Matlab.

Detailed descriptions of using an AOSLO for measuring visual sensitivity have been published previously (Harmening, Tuten, Roorda, & Sincich, 2014; Tuten, Harmening, Sabesan, Roorda, & Sincich, 2017; Tuten, Tiruveedhula, & Roorda, 2012). Prior to each session, mydriasis and cycloplegia were induced via instillation of 1% tropicamide and 2.5% phenylephrine ophthalmic solutions. Subjects used a bite bar to minimize shifts in pupil position during testing. Prior to each measurement block, three 40-frame retinal videos were acquired in parallel from the infrared and stimulation channels on the native frame grabber. The spherical focus of the AO system was set by the examiner to maximize the apparent sharpness of the cone mosaic in the stimulus channel image—presumably an objective indicator of stimulus light focus in the photoreceptor plane (**Figure 1B**). Transverse chromatic aberration (TCA) was computed from these videos by comparing the spatial offsets in retinal structure observed between the synchronized 795-and 550-nm video frames (Harmening, Tiruveedhula, Roorda, & Sincich, 2012). To achieve this, each video frame was first de-sinusoided and converted to the pixel scaling of the FPGA system using the image transformation obtained prior to testing (see above). Next, corresponding infrared and visible-wavelength frames were full-frame registered using a method based on the discrete Fourier transform (Guizar-Sicairos, Thurman, & Fienup, 2008). For a single 40-frame video, the TCA measurement was taken as the median x-and y-offset, in FPGA pixels, of the frame-by-frame registrations. In most cases, TCA measurements were repeated after each experimental session, and the overall TCA was taken as the mean of the pre-and post-session values. TCA measurements were used in offline analyses to determine the retinal locus targeted for testing (see below). Due to the high light levels required to capture retinal images with green light, threshold measurements commenced no sooner than 10 minutes after collecting the last TCA video, thus ensuring any photopigment bleached during imaging was sufficiently regenerated.

The relationship between stimulus size and detection threshold was assessed for three stimulus conditions. All testing was done at, or near, the subject’s central fovea. In Condition 1, ocular high-order aberrations were corrected in closed loop (7.75 mm pupil) and stimuli were delivered stabilized on the retina. The entrance pupil diameter in the 550-nm stimulus channel was 7.75 mm; in the diffraction-limited case, the central core of the corresponding point spread function would have a full-width at half-maximum of 0.24 arcmin, smaller than a foveal cone (∼0.5 arcmin). In this condition, stimuli were targeted to the subject’s preferred retinal locus of fixation (PRL). The PRL was determined prior to testing by recording a video of the subject maintaining fixation on a small flashing spot (550 nm) for 5 seconds, after which the retinal locations that sampled the fixation stimulus train were extracted; the PRL was taken as the median of these points. We note here that the PRL does not necessarily co-localize with the region of peak foveal cone density, but it is typically not displaced by more than a fraction of a degree (Putnam et al., 2005). Condition 1 minimizes retinal image blur from pre-retinal factors.

In Condition 2, ocular aberrations were compensated for as in Condition 1 but stimuli were allowed to drift naturally across the retina as the eye moved during fixation. This condition was akin to that used in Dalimier and Dainty (2010).

In Condition 3, a 3 mm aperture was placed in a pupil plane in the stimulus channel, and stimuli were delivered through the AOSLO system while the subject wore their habitual refractive correction. The entrance pupils in the imaging and wavefront sensing channels were unchanged. The intensity of the stimulus source was adjusted to equate the retinal irradiance with Conditions 1 and 2. The subject adjusted the defocus of the AO system manually to optimize the perceived sharpness of an 8.7 by 8.7 arcmin square-wave grating composed of 0.14 arcmin horizontal bars presented through the stimulus channel. All other system aberrations, along with the subject’s own high-order aberrations, were left uncorrected. Fixational eye movements could not be compensated in this condition due to the degradation in retinal image quality that results from leaving high-order aberrations uncorrected. To a first approximation, Condition 3 can be considered comparable to the experimental setup of Davila and Geisler (1991), where both normal optics and fixational eye movements affect the image incident on the retina. Spectral and irradiance parameters of the stimulus and background, however, were not matched to those used by Davila and Geisler.

For each of the three conditions, Ricco’s area was determined using the classic areal summation paradigm. Increment thresholds were measured for 10 circular stimuli (λ = 550 ± 15 nm) ranging in diameter from 0.43 to 9.25 arcmin (3 to 64 pixels in our AOSLO, where 415 pixels = 1 degree). Stimuli were presented against the raster-scanned background subtending approximately 1.25 by 1.25⁰ and comprising three wavelengths: (1) an infrared (λ = 848 nm) superluminescent diode (SLD; Superlum, Carrigtwohill, Ireland) used for wavefront sensing; (2) a near-infrared (λ = 795 nm) SLD used for retinal imaging (Superlum, Carrigtwohill, Ireland); and (3) a small amount of light at the stimulus wavelength passed by the AOM in its nominally-off state. The irradiances at the cornea were 6 µW, 30 µW, and 0.004 nW for the component wavelengths (848 nm, 795 nm, and 550 nm, respectively), resulting in a cumulative background luminance of ∼8 cd/m^2^. The maximum power of the 550 nm stimulus was 24.6 nW (828 cd/m^2^). Stimulus intensity was controlled by the AOM in linearized steps with 8-bit resolution.

In each block of trials, stimulus presentation was randomly interleaved, with detection thresholds for each spot size determined using 20-trial adaptive staircases guided by a yes-no response paradigm (Watson & Pelli, 1983). Measurement blocks were repeated three times per experimental condition, so that trials for each stimulus size were presented a total of 60 times. Each trial was initiated by the observer via button press, triggering the recording of a one-second retinal video during which the stimulus was presented for 187.5 ms (3 video frames). Stimulus delivery was encoded into stimulus video frames by placing a fiduciary digital marker at the image pixel corresponding to the center of the delivered stimulus. The placement of the digital marker takes into consideration the time at which the AOM was engaged and the size of the stimulus, thus providing a nominal localization of each delivered stimulus relative to the cone mosaic observable in the infrared image. Determining the veridical location of the delivered stimulus requires incorporating shifts between the imaging and stimulation wavelength induced by TCA; this correction was done during data analysis using the measurements described above. The subject indicated whether they detected the stimulus using a second button press.

### Data analysis: determining Ricco’s area and relating it to foveal anatomy

After the experiment, trial videos were stabilized using offline image registration tools, and the location of stimulus delivery relative to the cone mosaic was determined for each stimulus frame. For Condition 1, where stimuli were explicitly targeted to the subject’s PRL, trials on which the stimulus delivery marker fell outside of a 4.75 x 4.75’ square window, centered on the median delivery location of all trials, were excluded from subsequent analyses. For a trial to be considered valid, all three stimulus frames had to be delivered within the inclusion window. No data were excluded in Conditions 2 and 3 on the basis of delivery location or stimulus motion on the retina. For each stimulus, valid trials were fit with a logistic psychometric function and threshold, defined as the intensity that was detected on 78% of trials, was extracted. To determine Ricco’s area, threshold energy (i.e. threshold intensity multiplied by stimulus area) was plotted as a function of stimulus area on log-log axes and fit with a two-segment linear regression. The y-intercept of the first segment was allowed to vary while its slope was constrained to be zero. The slope and intercept of the second segment were allowed to vary. Ricco’s area of complete summation was taken as the stimulus area at which the two segments intersected.

Estimates of measurement precision for thresholds and Ricco’s area were obtained using bootstrapping. First, at each stimulus size, trial data were resampled randomly with replacement and psychometric functions were refitted. Ricco’s area was determined from the bootstrapped threshold energies as a function of stimulus area, as described above. This process was repeated 500 times; error bars throughout the manuscript span the central 90% of the bootstrapped parameter distributions (i.e. the 5% to 95% percentiles). Psychometric functions were fit using the Palamedes Toolbox (Prins & Kingdon, 2009) and spatial summation curves were fit using the Matlab routine “nlinfit”. A one-way analysis of variance (ANOVA) was conducted to assess whether Ricco’s areas depended on test condition, with p-values less than 0.05 considered statistically significant.

To compare our measurements of foveal Ricco’s areas to the underlying photoreceptor mosaic structure, we collected high-resolution videos at the fovea using denser pixel sampling (0.75 x 0.75⁰) in the native acquisition configuration of our AOSLO (Dubra & Sulai, 2011). These videos were registered and averaged using offline image processing tools (Dubra & Harvey, 2010), and then scaled and aligned manually to the images collected during psychophysical testing. Circles representing Ricco’s area were overlain on these high-resolution images at the median stimulus delivery location and the number of cones encompassed by each subject’s Ricco’s area was determined using custom cone counting software (Garrioch et al., 2012). Specifically, all cones residing within a 5-by-5 arcmin box, centered on the median stimulus delivery location, were selected manually, and the local angular cone density was computed. The number of cones falling within Ricco’s area was estimated by multiplying the summation area by the angular cone density and rounding to the nearest integer.

### Estimating the spatial summation area with a computational observer

To investigate whether a single post-receptoral summation unit could account for our data collected both with and without ocular aberrations, we used an open-source simulation platform (ISETBio; https://github.com/isetbio). We modelled the series of transformations a stimulus undergoes as it proceeds through the ocular media and triggers photoisomerizations in the cones. We then used the simulated photoisomerizations to train a computational observer and estimate psychophysical performance. The first stage of the model included a specification of the spectral and radiometric properties of the stimulus incident on the cornea. These values were set to match the wavelengths and corneal irradiances used in the study. Next, the retinal image irradiance was estimated by passing the stimulus representation through an optical model of the human eye. For Condition 1, the eye was modeled using diffraction-limited optics with an 8 mm pupil. To approximate Condition 3, the model eye’s point spread function (PSF) was computed using the mean values of the Zernike coefficients measured across a population of 100 subjects, with the computed PSF corresponding to a 3 mm pupil (Thibos, Hong, Bradley, & Cheng, 2002).

The retinal image was then sampled by a hexagonally packed cone mosaic (0.26 x 0.26 degrees, 635 cones, density 104,000 cones/mm^2^) and an L:M:S ratio of 0.67:0.33:0.0. Cones had a 3 µm inner segment aperture, 5 ms integration time, and the cone fundamentals were those of Stockman and Sharpe (Stockman & Sharpe, 2000). These cone fundamentals incorporate light absorption by the lens and macular pigment as well as the absorbance spectra of the cone photopigments. Photopigment optical density was taken as 0.5 (from http://www.cvrl.org) and isomerization quantal efficiency (fraction of quantal absorptions resulting in an isomerization) as 0.67 (Rodieck, 1998). Photoisomerization responses (number of isomerizations per each 5 ms integration time) were computed for each cone over a 155 ms window which included a 100 ms stimulus presentation window at a temporal resolution of 5 ms. The simulated stimulus duration was specified as shorter than our actual stimulus duration as a computationally convenient way to specify a rough total integration time of 100 ms for visual information. A total of 2,000 response instances for each cone in the mosaic were computed for each stimulus, with the individual instances differing by independently drawn Poisson isomerization noise. In these simulations there were no eye movements.

Computational observer psychometric functions (detection rate vs increment stimulus energy) for the case where there is no post-receptoral neural summation were computed for each stimulus size as follows. For each stimulus energy, isomerization response maps (635 cones x 31 time bins) to that stimulus, and to a zero stimulus energy case were concatenated, forming a vector with 39,370 entries (39,370 = 635 cones x 31 time bins x 2 intervals). This simulated a two-interval task. In half of the 2,000 trials, the stimulus response vector was inserted first and the zero-energy response vector was inserted second, and in the remaining trials, the ordering was reversed. A principal components analysis (PCA) on the set of 2,000 response vectors was conducted and the first 60 principal components were retained. The resulting data were used to train a binary linear support vector machine (SVM) classifier with 10-fold cross-validation to classify responses that had the stimulus response component first versus those which had the stimulus response component second. The out-of-sample (cross-validated) misclassification rate (*r_err_*) was computed, and the value of the psychometric function at the examined stimulus energy was defined as 1-*_rerr_*. The functions were fit with a smooth sigmoidal curve and thresholds were defined as the energy at which the psychometric function crossed 75%. Computational observer thresholds were determined for each of our experimental stimulus sizes, and for four additional stimulus sizes near our psychophysical estimate of Ricco’s area. The latter were inserted to better characterize the point at which our experimental threshold energy-vs-area curves begin to deviate from complete summation. The stimulus sizes used were: 0.43, 0.58, 0.87, 1.16, 1.35, 1.73, 2.00, 2.31, 2.70, 3.10, 3.47, 4.63, 6.94, and 9.25 arcmin^2^.

To model post-receptoral neural summation, computational observer psychometric functions for the same stimulus set were also computed with a post-receptoral summation stage. In this case, a Gaussian weighted pooling kernel, centered on the central cone in the simulated mosaic, integrated cone photoisomerizations over space. This reduced the number of entries in each simulated response vector to 62 (62 = 1 kernel x 31 time bins x 2 intervals). These data were also subjected to binary SVM classification (as described above but without PCA projection). Computational observer thresholds were computed using 6 log-spaced kernel sizes, with Gaussian standard deviations (*σ*) ranging from 0.125 to 4 arcmin (kernel areas 0.20 to 201 arcmin^2^, where the Gaussian radius = 2 *σ*).

As with our experimental data, these simulated threshold energies were plotted as a function of stimulus area. Qualitative inspection revealed that our experimental summation curves for both Conditions 1 and 3 were bracketed by the computational observer summation curves generated with σ = 1.0 and σ = 2.0 arcmin kernels. Because the simulations are computationally demanding, we estimated the intermediate kernel size that best accounted for our data by linearly interpolating between the two curves. Specifically, at each stimulus size, the interpolated log threshold energy, *T_interp_*, was computed using Equation 1:

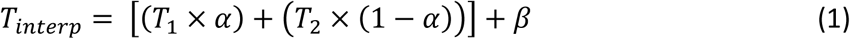

where *T_1_* and *T_2_* are the computational observer log threshold energies for the 1.0 and 2.0 arcmin kernels, respectively; *α* is a weighting term; and *β* is a vertical shift applied to the curve to account for absolute sensitivity differences between the computational observer and human subjects.

For each condition, average log threshold energies were computed by shifting each subject’s data set vertically to align its mean with the grand mean across subjects. Next, the parameters α and β were varied until the root-mean squared error between the experimental and model summation curves was minimized. The interpolated summation kernel size, *σ_interp_*, was computed using Equation 2:

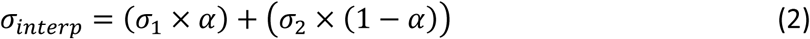

The summation kernels that best accounted for the experimental data were computed independently for Conditions 1 and 3. To examine whether a single post-receptoral summation filter of fixed spatial extent could account for the shape of the summation curve observed in both conditions, the data from both conditions were also fit simultaneously (i.e., *α* was constrained to be equal across conditions). In this case, *β* was determined independently for each condition. For comparison, threshold energy-versus-area curves were also generated in a variant of the model which included no post-receptoral summation using the approach described above.

## Results

Imaging the retina with an adaptive optics-equipped ophthalmoscope enables the visualization of individual photoreceptor cells in living eyes (Liang, Williams, & Miller, 1997; Morgan, 2016). In addition to its ability to reveal outer retinal structure with cellular resolution, the AOSLO used in this study confers two additional experimental advantages. First, a confocal light detection scheme is employed in each imaging channel, thereby enabling the simultaneous acquisition of full-frame images at different wavelengths with high axial resolution (**Figure 1A**). This capability affords a precise and objective verification of AO-correction fidelity for both the imaging and stimulation wavelengths based on the assumption that, for a given wavelength, an optimally-corrected eye will produce a clearly-resolved, high-contrast image of the cone mosaic. **Figure 1B** shows how introducing a small amount of defocus (±0.10 diopters [D]) into the 550 nm stimulus channel can degrade image quality appreciably. For comparison, psychophysical blur detection thresholds for foveal viewing are approximately ±0.20 D under AO-corrected conditions (Atchison & Guo, 2010; Atchison, Guo, Charman, & Fisher, 2009). Second, imaging the retina with a raster-scanning system permits the estimation of retinal motion at frequencies that exceed the nominal frame rate via strip-based image registration (Vogel et al., 2006); this motion signal can be harnessed to deliver stimuli in a retinally-contingent fashion with an accuracy on the order of 0.15 arcmin (Arathorn et al., 2007; Yang et al., 2010). Together, these features facilitate the experimental manipulations required to examine the contributions of optical aberrations and fixational eye movements to foveal spatial summation.

Spatial summation curves for AO-corrected, retinally-stabilized stimuli (Condition 1) delivered to the central fovea are shown in **Figure 2**. These plots depict threshold energy as a function of stimulus area; under a complete summation regime, the former is independent of the latter. For these conditions, the average diameter of complete summation (dashed black lines, **Figure 2**) extracted from the threshold energy-versus-area curves was 2.41 arcmin (range: 2.20 to 2.94 arcmin; see **Table 1**). The average slope of the second branch of the two-segment linear fit was 0.59 (range: 0.57 to 0.61). If these slopes were 1, it would indicate that stimulus energy falling outside of the summation area had no effect on threshold. The fact that the measured slopes are less than 1 but greater than 0 indicates that there is partial summation of this stimulus energy for spot sizes which exceed Ricco’s area.

**Figure 2.**
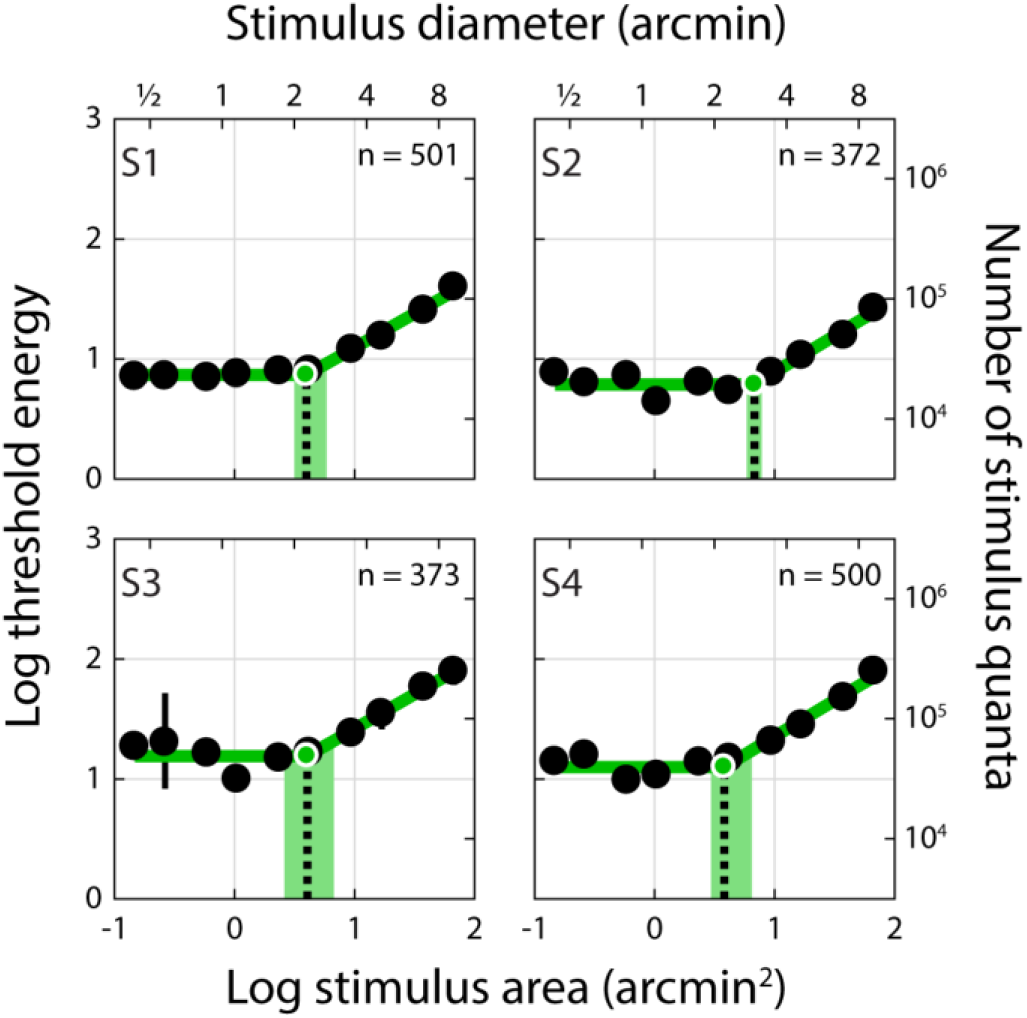
Threshold energy plotted against stimulus area for AO-corrected, retinally-stabilized stimuli delivered to the foveal center (Condition 1). Threshold energy-versus-area plots for Condition 1. Subject number is indicated in the upper left corner of each panel. Black dots represent increment threshold energy for each stimulus size. Thresholds energy units follow the convention of Davila and Geisler (1991): threshold luminance (cd/m^2^) x stimulus area (arcmin^2^) x stimulus duration (seconds). Threshold error bars span the 5^th^ and 95^th^ percentiles of the distribution obtained from the bootstrapping procedure; where no error bars are shown, this range is smaller than the plotted symbol. Stimulus diameter is provided on the secondary x-axis; threshold energy expressed as number of increment stimulus quanta incident on the cornea is provided on the secondary y-axis. The green line shows the two-segment linear regression, where the slope of the first branch was constrained to be zero (i.e. complete summation); Ricco’s area (black dashed line) was taken as the intersection of the two-segment fit. The green shaded area spans the 5^th^ to 95^th^ percentiles of the bootstrapped Ricco’s area distribution (see Methods). The number of trials (out of 600) satisfying the stimulus delivery criterion for inclusion in this analysis is indicated in the upper right corner of each panel.

**Table 1.** Summary of summation areas for all subjects and conditions.

| Subject | Summation areas in arcmin<sup>2</sup><br>(5<sup>th</sup> – 95<sup>th</sup> bootstrap percentiles) |  |  |
| --- | --- | --- | --- |
|  | Condition 1<br>AO, retinally-stabilized | Condition 2<br>AO, with FEM | Condition 3<br>No AO, with FEM |
| S1 | 3.97<br>(3.16 – 5.83) | 4.12<br>(3.33 – 6.92) | 6.17<br>(3.92 – 7.53) |
| S2 | 6.80<br>(5.85 – 7.78) | 7.26<br>(4.18 – 8.43) | 6.85<br>(5.51 – 8.40) |
| S3 | 4.04<br>(2.60 – 6.70) | 6.28<br>(3.58 – 8.42) | 6.20<br>(3.71 – 7.15) |
| S4 | 3.80<br>(2.95 – 6.43) | 6.85<br>(3.99 – 7.94) | 8.28<br>(3.92 – 16.81) |
| Average (± SD) | 4.65 ± 1.44 | 6.13 ± 1.40 | 6.88 ± 0.99 |
| Davila & Geisler (1991)* | — | — | 5.48 ± 2.07 |
| Dalimier & Dainty (2010)<sup>†</sup> | — | 5.03 ± 2.78 | — |

\*3 mm pupil with natural optics; 8-10 cd/m<sup>2</sup> background; n = 4 subjects
<sup>†</sup>6 mm pupil with AO correction; 20 cd/m<sup>2</sup> background; n = 3 subjects

The summation diameters we measured were about 5 times larger than the inner segment diameter of a foveal cone in the human retina (∼0.5 arcmin; Curcio et al., 1990; Hirsch & Curcio, 1989), suggesting that our stimulus engaged detection mechanisms that pool photons completely across multiple foveal cones. The imaging capabilities of the AOSLO permit a direct comparison of the summation areas we obtained in Condition 1 with the structure of the foveal cone mosaic on a subject-by-subject basis. In **Figure 3**, each panel depicts an image of a subject’s fovea with a circle representing Ricco’s area placed at the median stimulus delivery location. These images, which were acquired over a smaller field of view than those acquired during psychophysical testing to allow for finer pixel sampling, show that cones within our subjects’ foveas could be resolved and counted directly. The number of cones comprising Ricco’s area were 20, 37, 23, and 17 in our four subjects (S1-S4, respectively).

**Figure 3.**
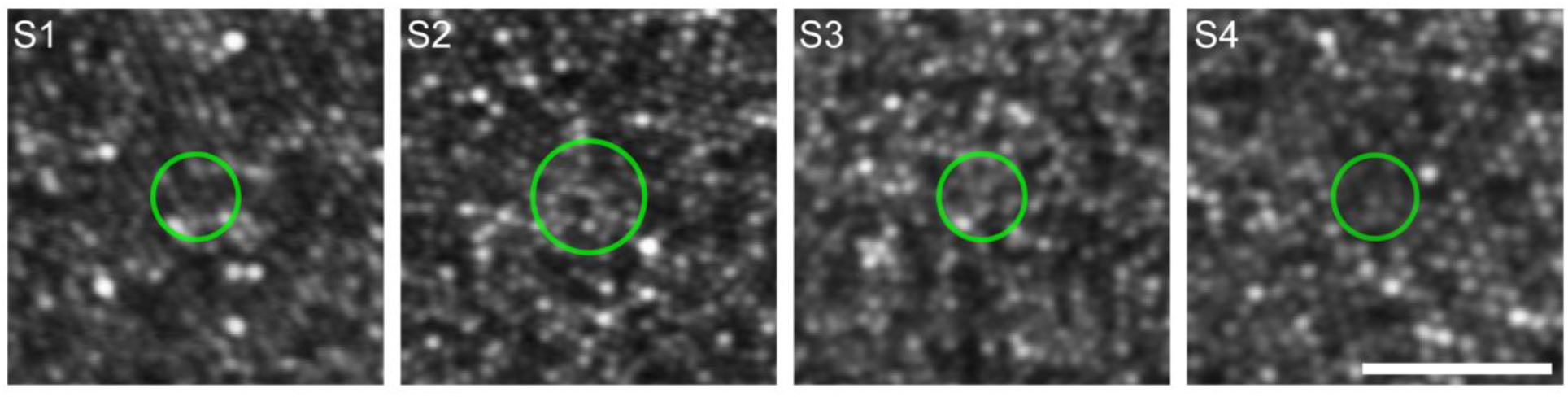
Foveal summation areas from Condition 1 compared to the underlying cone mosaic. High-resolution retinal images from Subjects 1 through 4 show densely-packed cone photoreceptors in the foveal region. Each image was generated by averaging several spatially-registered AOSLO video frames (number of frames, from left to right: 50, 30, 45, and 45). Green circles represent Ricco’s area of complete summation obtained for each subject in Condition 1; summation markers are placed at the median stimulus delivery location on the retina after accounting for the effects of transverse chromatic aberration. Scale bar represents 5 arcmin.

In Condition 2, stimuli were allowed to drift naturally across the retina due to fixational eye movements. Using this approach, we found that on average Ricco’s diameter was 2.78 arcmin (**Figure 4;** range: 2.29 to 3.04 arcmin). These values are similar to those obtained by Dalimier and Dainty (2010) using an AO vision simulator (**Table 1**). Although relative to Condition 1, each subject’s Ricco’s area increased slightly in Condition 2, the respective parameter distributions obtained via bootstrapping overlapped substantially (**Table 1**). One potential explanation for the similar summation measurements we observed in Conditions 1 and 2 is that the magnitude of stimulus motion on the retina was not significantly different between the two conditions. To investigate this possibility, we examined the retinal videos acquired during each trial, extracted the delivered location on the retina, and computed the linear distance traversed by the stimulus over the three-frame (187.5 ms) presentation epoch. Trials featuring microsaccades, blinks, or diminished image quality—all of which undermine the image registration necessary for accurate determination of stimulus trajectories—were excluded from this analysis. When fixational eye movements were compensated for by the eye tracking software (Condition 1), the median stimulus travel averaged 0.58 arcmin (range: 0.47 to 0.64 arcmin; **Figure 5**, green histograms) across our four subjects, approximately equal to the angular subtense of a single foveal cone. For Condition 2, involuntary fixational eye motion produced a roughly threefold increase in intratrial stimulus motion, with an average (across subjects) median of 1.79 arcmin (range: 1.56 to 2.18 arcmin; **Figure 5**, gray histograms). The drift amplitudes we observed were in line with those reported previously for similar temporal intervals (Cherici, Kuang, Poletti, & Rucci, 2012). When Ricco’s areas were recomputed using only the Condition 2 trials included in this analysis, they did not differ significantly (p = 0.875; Wilcoxon signed-rank test; average diameter: 2.79 arcmin; range: 2.27 to 3.08 arcmin) from those obtained using all Condition 2 trials (i.e. the data shown in **Figure 4**). Ricco’s areas for Condition 1 were not recomputed, because the inclusion criterion used here was the same as that incorporated into the analyses behind **Figure 2**.

**Figure 4.**
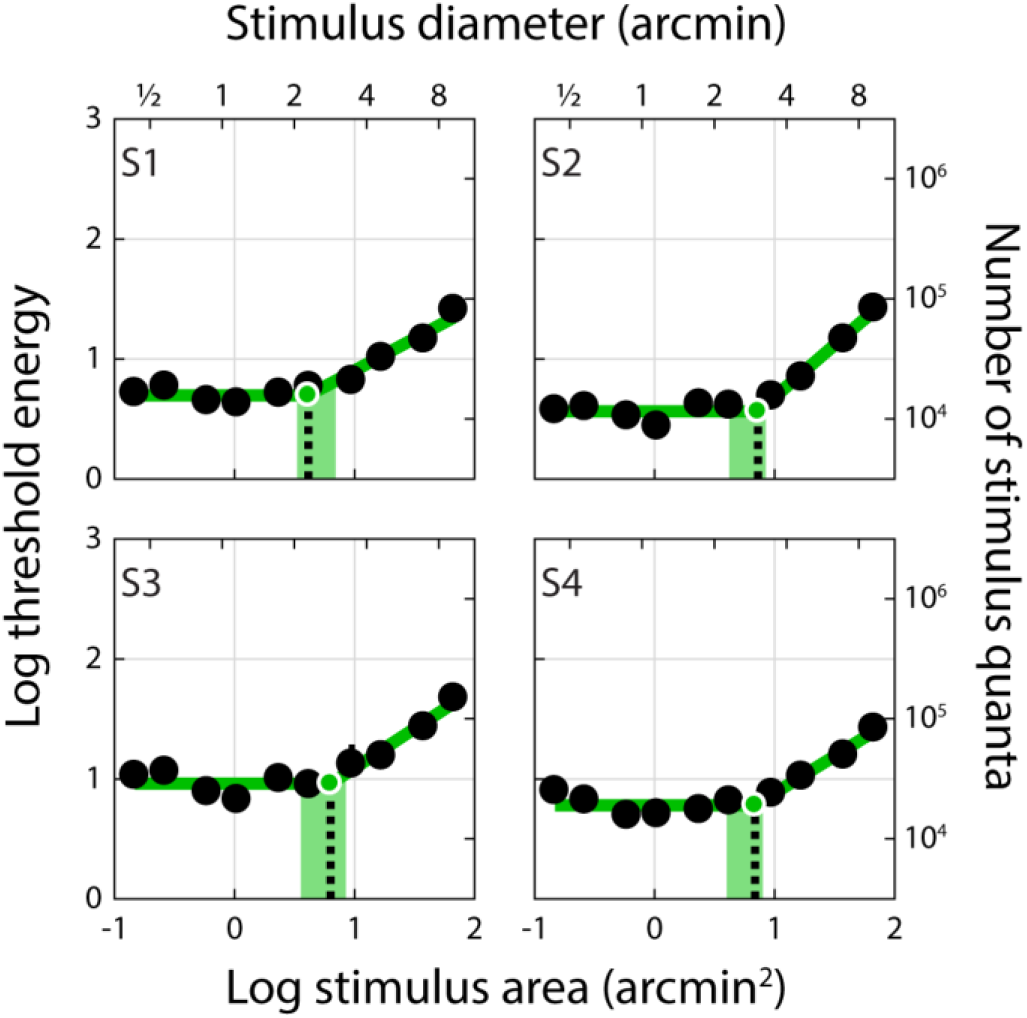
Threshold energy plotted against stimulus area for AO-corrected, non-stabilized stimuli delivered to the fovea (Condition 2). Summation curves for Condition 2. All else as in Figure 2.

**Figure 5.**
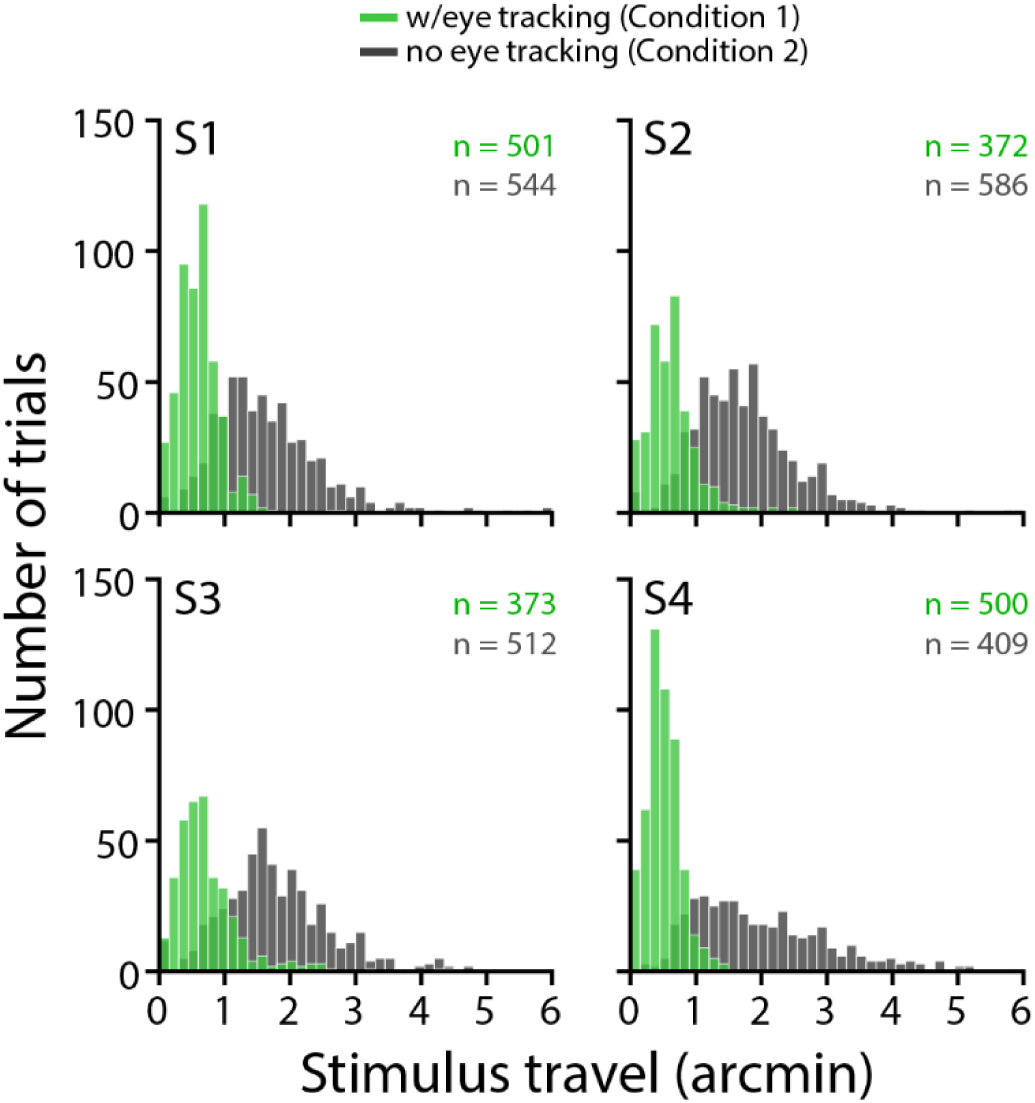
Intra-trial stimulus travel on the retina for Conditions 1 and 2. Histograms of the total angular distance traversed by the stimulus on the retina during each 3-frame presentation. Green histograms show the distribution of stimulus motion magnitude with eye tracking and retinally-contingent delivery (Condition 1), while gray bars depict stimulus motion that resulted when the stimulus was allowed to drift naturally across the retina as the eye moved during the presentation interval (Condition 2). Each panel corresponds to a single observer, with the number of included trials (out of 600) shown in the upper right corner (green text = Condition 1; gray text = Condition 2). Condition 1 trials shown here correspond to those included in the summation curves presented in Figure 2. For Condition 2, trials were excluded from this analysis when the image registration required to compute stimulus trajectories was corrupted by microsaccades, blinks, or diminished image quality; however, we note these trials were not excluded from the plots shown in Figure 4. Ricco’s areas computed with the subset of Condition 2 trials included in this analysis were statistically indistinguishable from those computed from all trials (see Results).

Summation curves for Condition 3 are shown in **Figure 6**. These measurements were obtained with stimuli projected through the AOSLO with the subject wearing their habitual refractive correction and a stimulus-channel pupil size of 3 mm. Despite not compensating for ocular aberrations or fixational eye movements, we obtained summation curves similar to Conditions 1 and 2, with an average Ricco’s diameter of 2.95 arcmin (range: 2.80 to 3.25 arcmin). Individual summation areas for Conditions 1 through 3, along with summary results from Davila and Geisler (1991) and Dalimier and Dainty (2010), can be found in **Table 1** and **Figure 7**. Summation areas did not change significantly with test condition (one-way ANOVA; F_2,9_ = 3.08; p = 0.096).

**Figure 6.**
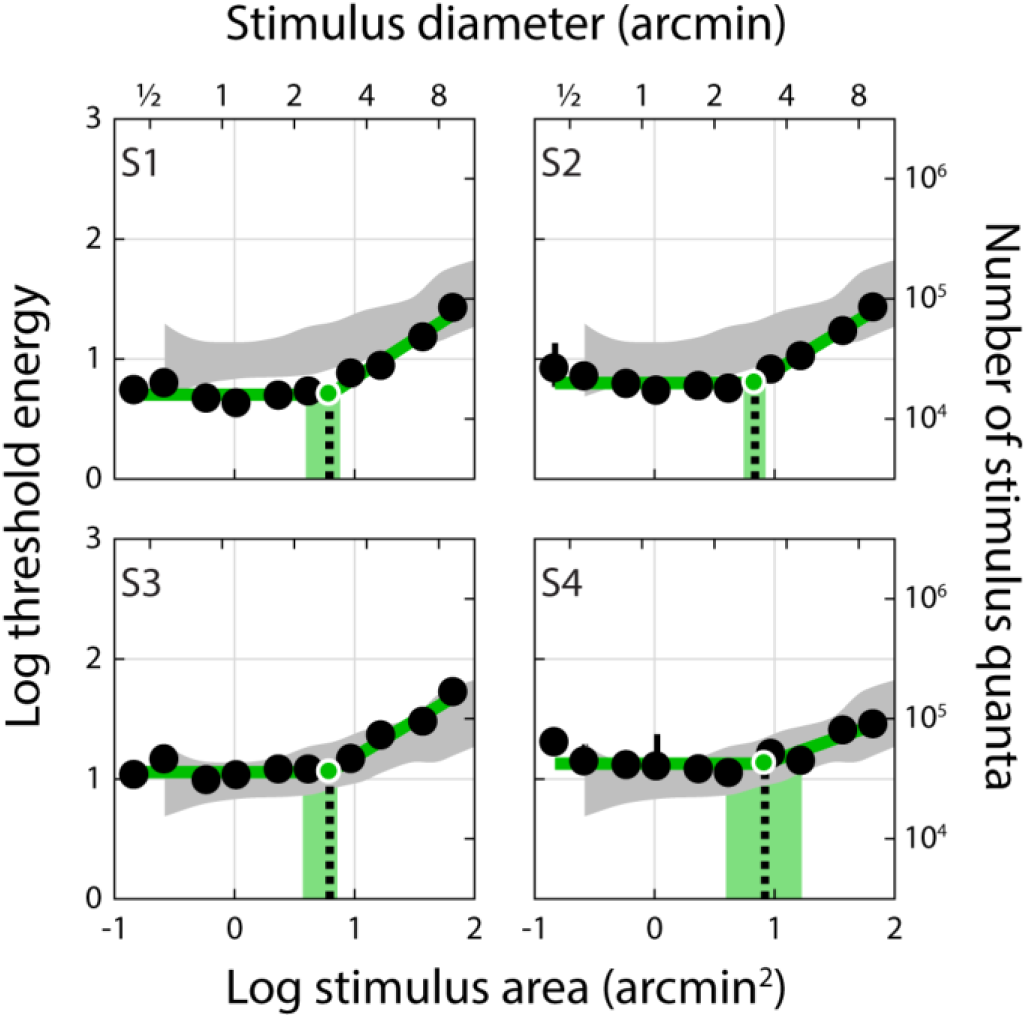
Threshold energy plotted against stimulus area for non-stabilized, natural optics stimuli (3 mm pupil) delivered to the fovea (Condition 3). Summation curves for Condition 3. Gray shaded regions show data (mean ±2 SD) from Davila and Geisler (1991); these data were shifted down 0.09 log units to account for the slight difference in background luminance between their study (10 cd/m^2^) and ours (8 cd/m^2^), presuming a Weber adaptation regime. No adjustment was made for the difference in wavelength composition of the stimuli between our experiment and those of Davila and Geisler. All else as in Figure 2.

**Figure 7.**
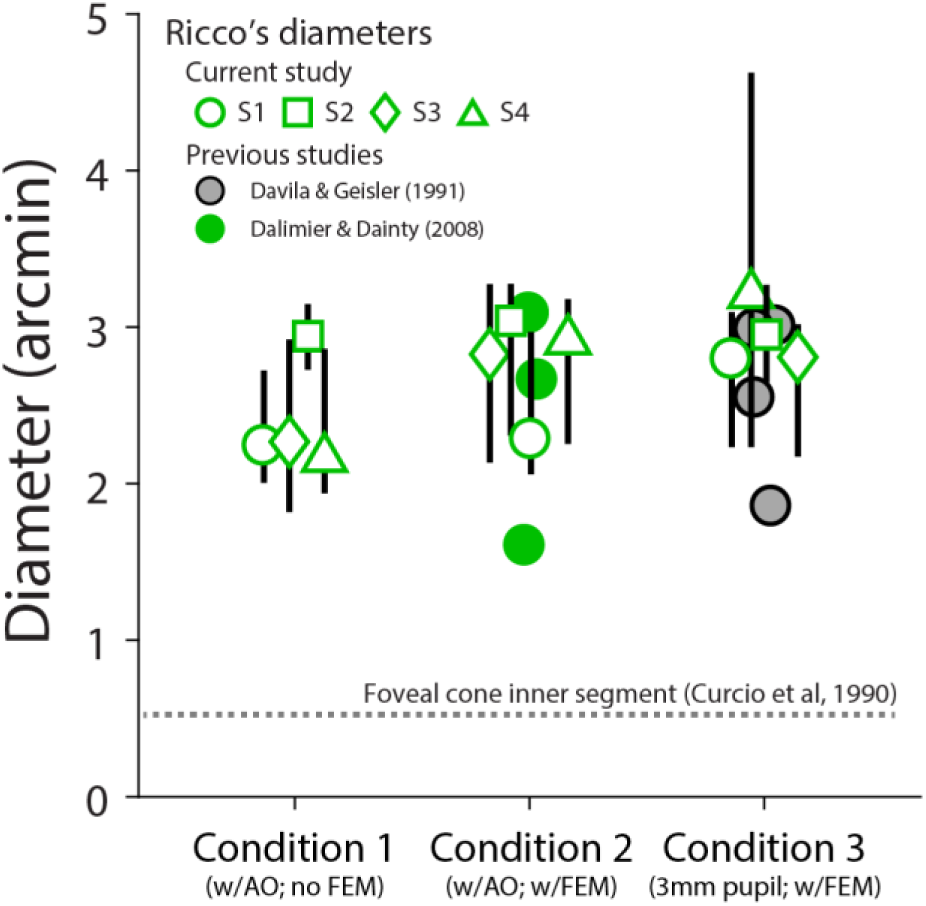
Ricco’s diameters for Conditions 1 through 3 compared to previous studies. Ricco’s diameters (in arcmin) are plotted as open green symbols for each experimental condition in the present study. Subject is indicated by marker shape. Data points are jittered horizontally to enhance visualization. Error bars span the central 90% of the bootstrapped Ricco’s diameter parameter distributions (see Methods). Individual data points from previous studies with similar experimental conditions are shown for comparison. Gray dashed line represents the angular diameter of a foveal cone in the human retina (Curcio et al., 1990).

To assess whether our data collected both with and without correction for ocular aberrations are consistent with a single post-receptoral summation unit, we compared the threshold energy-versus-area curves we obtained in Conditions 1 and 3 with simulations generated using a computational observer with diffraction-limited and natural optics, respectively. A schematic of the model architecture is shown in **Figure 8A**. The average threshold energies across subjects for Condition 1 are plotted as a function of stimulus area in the top panel of **Figure 8B**. The flat branch on the left side of this curve is indicative of complete summation of stimulus energy, while the rising branch results from partial summation as stimulus size increases. We first simulated summation curves using diffraction-limited optics without a post-receptoral pooling stage, the result of which was a monotonically-increasing function (**Figure 8B**, top panel, gray line); the faint flattening at the smallest stimulus sizes corresponds to summation across the cone aperture (MacLeod, Williams, & Makous, 1992). Introducing a post-receptoral pooling kernel to the simulation produces characteristic summation appearance in the left side of the threshold energy-versus-area curve (**Figure 8B**, top panel, black line). The summation kernel sigma that best fit our Condition 1 data was 1.6 arcmin, corresponding to a full-width at half-maximum (FWHM) of 3.77 arcmin. As one would expect, this is larger than the mean Ricco’s area extracted from fitting the same data with a two-segment linear regression, because the latter assumes perfect summation rather than Gaussian-weighted summation.

**Figure 8.**
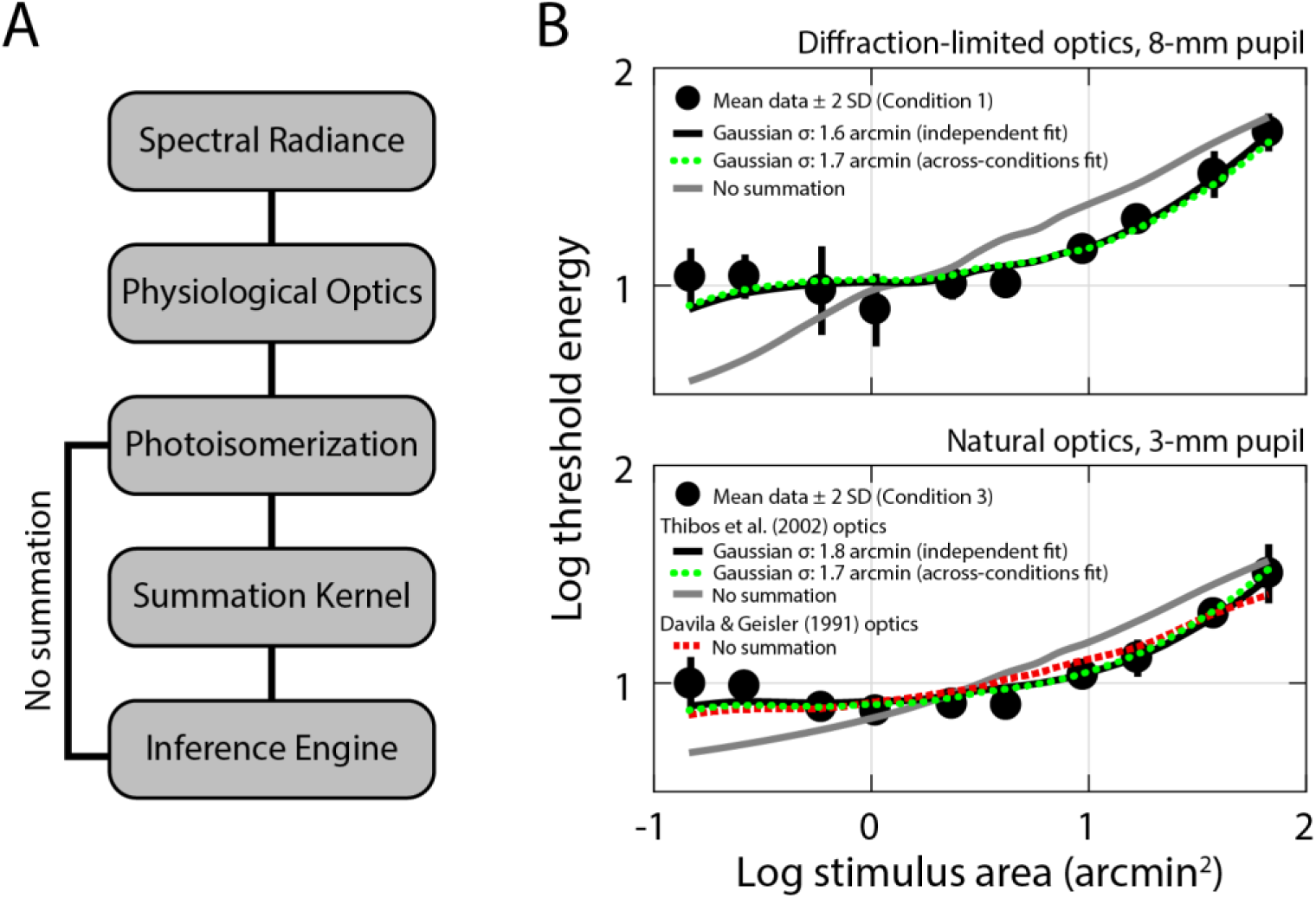
Modeling summation curves with a computational observer. (A) Schematic of computational observer stages; see Methods for details. (B) Mean threshold energies (black dots) from Condition 1 (top panel) and Condition 3 (bottom panel) are plotted as a function of stimulus area. Prior to averaging, threshold energy data for each subject were shifted vertically to bring the intra-subject mean into alignment with the grand mean for all subjects. Error bars are ± 2 SD. Simulated summation curves generated with the computational observer are also shown; all simulations in the top panel were generated using a computational observer featuring diffraction-limited optics, while those in the bottom panel were obtained when a standard model of ocular aberrations was incorporated into the computation (see Methods). Fixational eye movements were not incorporated into the computational observer. The solid black lines denote the simulated summation curves generated with post-receptoral summation kernels that best fit each condition independently, whereas the dashed green lines show the simulations that result when the summation kernel was constrained to be the same across conditions. The solid gray lines show simulations from a computational observer with no post-receptoral summation using the population mean for optical aberrations reported in Thibos et al. (2002); the red dashed line in the bottom panel shows the simulation generated using the optical model specified in Davila and Geisler (1991). All simulation curves were allowed to shift vertically as part of the fitting procedure (see Methods).

Mean threshold energies for Condition 3 are plotted in the bottom panel of **Figure 8B**, along with simulations obtained with a computational observer equipped with typical ocular aberrations (Thibos et al., 2002). Our Condition 3 data were best matched by simulated data (**Figure 8B**, bottom panel, black line) generated with a post-receptoral pooling kernel sigma of 1.8 arcmin (FWHM = 4.24 arcmin). When the respective simulations were fit simultaneously to data from Conditions 1 and 3, the optimal kernel sigma was 1.7 arcmin (FHWM = 4.00 arcmin), producing similar simulated curves (**Figure 8B**, green dashed lines) to those generated independently. By contrast, the computational observer simulation (gray line) without a post-receptoral pooling stage demonstrates that although typical amounts of optical aberrations (Thibos et al., 2002) can produce a summation-like appearance, the extent of spatial pooling due to optical factors was insufficient to capture the shape of our data. We also confirmed we could replicate the core result from Davila and Geisler (1991): when the computational observer simulation was repeated using the same optical model that they had available (derived from the line-spread measurements of Campbell and Gubisch (1966)), the summation arising from optical spread closely resembled our Condition 3 data (**Figure 8B**, bottom panel, red dashed line). Across all simulations, the vertical shifts applied to align the computational observer data with their psychophysical counterparts ranged between 2.3 and 2.5 log units.

## Discussion

The degree to which optics, eye motion, photoreceptor sampling, and post-receptoral neural processing combine to shape visual performance is a longstanding and fundamental question in vision science. We used an adaptive optics system in conjunction with precise retinal tracking to measure spatial summation in the human fovea. When high-order aberrations and stimulus motion on the retina were minimized experimentally, we measured Ricco’s areas in the central fovea that exceeded the dimensions of a single foveal cone (**Figures 2** and **3**). When we repeated our measurements with ordinary levels of optical aberrations and stimulus motion (**Figures 4** and **6;** see also **Table 1** and **Figure 7**), the summation areas did not change significantly, and were generally consistent with previous psychophysical estimates of spatial pooling in the mechanisms mediating foveal contrast detection (Dalimier & Dainty, 2010; Davila & Geisler, 1991; Levi & Klein, 1990). Our results demonstrate that the pooling of individual cone signals by post-receptoral circuitry plays an important role in spatial summation in the fovea.

The present study builds upon previous attempts to parse the relative contributions of pre-neural factors to spatial summation in the fovea. Davila and Geisler (1991) conducted an ideal observer analysis in which summation curves were generated using a model of the early visual system that incorporated contemporary estimates of the optical quality of the eye and the arrangement and quantum sensitivity of the cone mosaic. They concluded that optical spread produced by the refractive components of the eye was sufficient to replicate the degree of summation they observed psychophysically. While Davila and Geisler’s model did not explicitly require additional neural summation in the cone-mediated pathway, nor could the possibility of post-receptoral pooling on a scale commensurate to optical blurring be excluded by their calculations. Disentangling the two sources of spatial summation requires experimental control of ocular aberrations, a capability not realized until the advent of AO for studying human vision (Liang et al., 1997). Indeed, a computational observer without post-receptoral summation provides a reasonable fit to our Condition 3 data when the optical model of Davila and Geisler is used (**Figure 8B**, bottom panel, red dashed line). However, measurements obtained when pre-retinal factors were minimized (**Figure 2**) resolves this ambiguity, and provides strong evidence for the existence of mechanisms in the foveal circuitry that pool signals across multiple cones at detection threshold.

More recently, Dalimier and Dainty (2010) reported a significant reduction in Ricco’s area when high-order aberrations were minimized with an AO vision simulator, although a 6 mm pupil was used in both conditions. By contrast, with a 3 mm pupil and natural optics, we obtained spatial summation curves which were essentially indistinguishable from those measured with AO correction over a 7.75 mm pupil (**Figure 7**). Thus, it appears that high-order ocular aberrations do not contribute to foveal summation at pupil sizes commonly observed at photopic light levels (Winn, Whitaker, Elliott, & Phillips, 1994).

The summation areas we measured when pre-retinal factors were minimized with AO encompassed, on average, roughly two dozen foveal cones. This level of pooling far exceeds the amount presumed to exist in the fine-grained parvocellular retinogeniculate pathway, suggesting that the anatomical basis of Ricco’s area may reside elsewhere. Previous investigators have proposed that Ricco’s area is correlated with anatomical features of parasol retinal ganglion cells (Volbrecht et al., 2000), neurons which project to the visual cortex via the magnocellular pathway. In macaque retina, histological evidence suggests parasol cells near the fovea draw excitatory input from 30 to 50 cones (Calkins & Sterling, 2007; Grünert, Greferath, Boycott, & Wässle, 1993). Although these numbers are similar to the number of receptors we estimated to underlie Ricco’s area in our subjects (**Figure 3**), parasol dendritic arbor diameters in the human fovea are thought to be about twice as wide as their macaque counterparts: angular diameters of approximately 7 arcmin have been estimated from a handful of cells located ∼0.5 degrees from the foveal center (Dacey & Petersen, 1992; Kaplan, Lee, & Shapley, 1990). If similar dimensions are maintained in parasol cells sampling the foveola, they would be driven by anywhere from 120 to 200 cones, depending on the local photoreceptor packing density (Curcio et al., 1990; Goodchild, Ghosh, & Martin, 1996). If this were the case, our measurements may not reflect complete signal integration across the entire parasol dendritic field, but rather arise from complete summation occurring within neural units of intermediate size.

One alternative retinal substrate for spatial summation within the magnocellular pathway is diffuse bipolar cells, interneurons that relay cone signals to parasol ganglion cells. In various species and ganglion cell classes, bipolar cells serve as receptive field subunits that sum cone inputs linearly (Freeman et al., 2015; Liu et al., 2017; Schwartz et al., 2012) and introduce a rectifying nonlinearity into the signal transfer at the bipolar-ganglion cell synapse (Demb, Haarsma, Freed, & Sterling, 1999; Demb, Zaghloul, Haarsma, & Sterling, 2001). The electrophysiological signature of this nonlinear summation is a frequency-doubling response to counterphase-modulated gratings, a phenomenon first observed in recordings of cat retinal ganglion cells (Enroth-Cugell & Robson, 1966; Hochstein & Shapley, 1976a, 1976b). More recently, nonlinear spatial summation has been revealed in primate parasol ganglion cells, implying that their receptive field centers may also feature some level of subunit organization (Crook et al., 2008; Petrusca et al., 2007). In the macaque retina, the convergence of cones onto diffuse bipolar cells is largely invariant with eccentricity: each neuron draws input from between 5 and 10 underlying photoreceptors (Boycott & Wässle, 1991; Grünert, Martin, & Wässle, 1994). Although these numbers are too low to account for the extent of summation in our data (roughly 20-40 cones; **Figure 3**), the discrepancy could be reconciled if diffuse bipolar cells in the human fovea exhibited the same fourfold increase in dendritic field area (relative to macaque) that has been reported for foveal parasol ganglion cells (Dacey & Petersen, 1992). In any case, further elucidation of the precise anatomy and physiology of visual pathways originating at the foveal center will be required before the neurobiological underpinnings of Ricco’s area can be determined with confidence.

Prior studies have shown the influence optical factors have on visual performance varies by task. For example, in the fovea, visual acuity and high-spatial frequency contrast sensitivity improve with defocus correction (Campbell & Green, 1965) and compensation of high-order aberrations (Rossi, Weiser, Tarrant, & Roorda, 2007; Yoon & Williams, 2002), reflecting their presumed reliance on a parvocellular substrate equipped with a “private-line” wiring scheme. However, if the Ricco’s areas we measured in this study are determined by pooling within a coarser neural pathway, it seems reasonable that performance on our task would be robust to modest amounts of blur, provided the spatial spread introduced by the defocus does not exceed the dimensions of the summation zone. While beyond the scope of the present study, the deleterious role of blur, as well as evidence of finer-grained neural processing, may become evident if summation measurements were repeated under conditions which favor detection by the parvocellular pathway (Pokorny & Smith, 1997; Smith, Sun, & Pokorny, 2001).

Our results also demonstrate that ordinary levels of fixational eye movements do not exert a meaningful influence on psychophysical measurements of post-receptoral pooling in the fovea. This outcome could be attributed to the similarity between the spatial extent of the summation areas we measured (∼2.5 arcmin diameter; **Table 1**) and the stimulus motion magnitudes we observed when eye movements were not compensated (∼1.79 arcmin; **Figure 5**). It is possible that larger summation areas could result from higher levels of stimulus motion. However, visual acuity and contrast sensitivity to high-frequency interference fringes—tasks reliant on mechanisms with even finer neural sampling—are generally unaffected when probed with a moving stimulus (Packer & Williams, 1992; Westheimer & McKee, 1975). Moreover, it has been suggested that the visual system may harness the spatiotemporal fluctuations in cone signals produced by eye movements to improve the detection of fine-grained targets (Kuang, Poletti, Victor, & Rucci, 2012; Ratnam, Domdei, Harmening, & Roorda, 2017; Rucci, Iovin, Poletti, & Santini, 2007; Rucci & Victor, 2015). The present findings are consistent with the view that the visual system is equipped with mechanisms capable of disregarding—and in some cases capitalizing on—the retinal image blur introduced by the unsteady eye.

Although our data obtained with and without AO correction could be accounted for by a relatively simple model incorporating a single post-receptoral summation stage with a FWHM of ∼4 arcmin (**Figure 8**), we do not conclude that the aggregate shape of the summation curve is determined solely by the activity of a single, univariant mechanism. The signals transduced in each cone are partitioned by the retina into at least 20 parallel retinogeniculate pathways (Dacey, Peterson, Robinson, & Gamlin, 2003), each of which tiles the retina and presumably transmits useful information about the visual scene that is then reassembled into a coherent percept at higher visual areas. The relative activity of these diverse pathways is likely stimulus-dependent. In an example relevant to the present study, multi-electrode array recordings in the peripheral retina have shown that single-cone modulations can drive midget and parasol cells with similar efficacy (Li et al., 2014); the superior contrast sensitivity traditionally attributed to the magnocellular pathway for larger stimuli appears to arise from pooling signals from multiple cones. From these results, it is conceivable that thresholds for the cone-sized spots in our paradigm could be determined by some mixture of midget and parasol ganglion cell activity, whereas detection of slightly larger circular increments may be mediated primarily by the latter (Swanson et al., 2011). A multiple-mechanism conception of spatial summation has been described previously using a cortical framework in which summation curves arise from pooling across a range of orientation-tuned spatial filters (Pan & Swanson, 2006). In such a scheme, equal-energy increment stimuli along the linear portion of the summation curve, though equally detectable, may nonetheless be discriminable along some other perceptual dimension, such as hue or apparent size.

## Acknowledgments

Supported by a Research to Prevent Blindness Stein Innovation Award, NIH U01EY025477, NIH R01EY023591, NIH P30 EY001583, Foundation Fighting Blindness, F. M. Kirby Foundation, Paul and Evanina Mackall Foundation Trust, Simons Foundation Collaboration on the Global Brain Grant 324759. B. A. Wandell, H. Jiang and J. E. Farrell contributed to the development of the ISETBio software used to develop the model.

